# Hopx expression marks Aβ clearance astrocytes in Alzheimer’s disease

**DOI:** 10.64898/2026.02.27.708674

**Authors:** Dong-Dong Cao, Jian Dong, Xiao-Yu Liao, Yang-Yang Teng, Yang Gu, Zhen-Ge Luo

**Affiliations:** School of Life Science and Technology, ShanghaiTech University, Shanghai, 201210, China; State Key Laboratory of Advanced Medical Materials and Devices, ShanghaiTech University, Shanghai, 201210, China

**Keywords:** Astrocytes, Aβ plaque phagocytosis, Astrocytic dysfunction, Hopx, Alzheimer’s disease

## Abstract

Astrocytic dysfunction is closely associated with nearly all types of neurological diseases. Targeting astrocyte regulation has thus emerged as an important potential therapeutic strategy for neurological disorders. However, mechanisms that induce astrocyte dysfunction under pathological conditions have not been fully elucidated. In this study, we identified that Homeodomain-only protein X (*Hopx*) is downregulated in astrocytes across inflammatory models, Alzheimer’s disease (AD) mouse models, and brain tissues from AD patients. In the brains of AD mice, Hopx-positive astrocytes exhibit efficient β-amyloid (Aβ) plaque clearance, while knockout of *Hopx* leads to astrocytic dysfunction. Conversely, astrocyte-specific overexpression of Hopx not only significantly enhances their Aβ phagocytic function but also effectively reduces the generation of neurotoxic astrocytes while increasing protective astrocytes. In summary, this study demonstrates the core regulatory mechanism underlying astrocytic dysfunction under AD pathological conditions and provides important potential targets for developing therapeutic strategies for AD by targeting astrocyte regulatory pathways.

## INTRODUCTION

Astrocytes are essential glial cells that play pivotal roles in maintaining central nervous system homeostasis^1, 2^. Under physiological conditions, they support neuronal function by regulating synaptic transmission, ion and neurotransmitter balance, metabolic supply, and blood-brain barrier (BBB) integrity^3–5^. In neuroinflammation and neurodegenerative disorders such as Alzheimer’s disease (AD), astrocytes undergo complex phenotypic and functional alterations that contribute to disease initiation and progression^6–8^. These changes can range from a loss of homeostatic support to the acquisition of neurotoxic properties, thereby promoting pathogenesis^9–11^. While reactive astrocytes facilitate the removal of toxic proteins, such as Aβ, they display neuroprotective effects and thereby mitigate disease progression^11–13^. Given the intricate and paradoxical nature of astrocytes under AD conditions, identifying the regulatory factors that govern their functional dynamics and subpopulation transitions presents a formidable challenge.

The transition of astrocytes from homeostatic to reactive states is governed by specific molecular orchestrators that sense pathological cues and drive transcriptional reprogramming, shifting astrocyte function along a spectrum from neuroprotective to neurotoxic phenotypes^13–17^. Recently, several studies using single-cell sequencing technology have comprehensively described changes in astrocyte subpopulations and their molecular expression profiles in AD mouse models and patients and under inflammatory conditions^8,14,18,19^. However, the intrinsic regulators that induce the neurotoxic state and regulate Aβ clearance under AD conditions remain poorly defined. By integrating bulk RNA-seq data of astrocytes from LPS-treated mouse^8^ and the single-cell RNA sequencing (scRNA-seq) data from 5×FAD mouse model^18^ and single-nucleus RNA sequencing (snRNA-seq) data from AD patients^14^, we identified the Homeodomain-only protein X (*Hopx*) as a candidate regulator consistently downregulated across all three datasets.

Hopx is a non-DNA binding homeodomain protein that acts as a regulatory scaffold, interacting with transcription factors and chromatin modifiers such as SRF, GATA factors, SMADs, and HDACs to regulate gene expression during heart development^20–22^. In the nervous system, Hopx is expressed in neural stem cells located in the subventricular zone (SVZ) and dentate gyrus (DG), as well as in astrocytes^23–25^. However, the role of Hopx in regulating astrocyte functions and astrocyte subpopulation transition has not yet been reported. Here, we found that Hopx-positive astrocytes exhibit increased Aβ phagocytic capacity, while ablation of these astrocytes exacerbates AD pathological features. Overexpression of Hopx in astrocytes enhanced clearance of Aβ plaques and promoted the proportion of protective astrocytes while decreasing the proportion of toxic ones. Overall, these findings demonstrate that astrocytic Hopx boosts astrocyte ability to eliminate toxic proteins and assists in restoring their normal astrocytic functions in AD pathogenesis, offering perspectives on the development of treatment strategies.

## RESULTS

### Downregulation of *Hopx* in murine and human astrocytes under AD conditions

Astrocyte reactivity is a complex and highly heterogeneous response to pathology, yet certain homogeneous substates persist across diverse disease conditions. For example, it has been shown that astrocytes adopt neuroprotective or neurotoxic substates in response to neuroinflammation, with similar transitions observed in neurodegenerative diseases^26, 27^. The nodal regulators controlling these transitions, however, remain poorly defined. To identify regulators responsive to multiple pathological stimuli, we analyzed differentially expressed genes (DEGs) from three contexts: disease-associated astrocytes (DAAs) versus Gfap-low astrocytes from AD mice^18^ (DAAs were reported as a population exhibited in AD mice but not in wild-type (WT) mice and Gfap-low astrocytes expressed hemostatic genes.), astrocytes from LPS-stimulated versus control mice^8^, and astrocytes from AD patients versus healthy controls^14^. While no genes were universally upregulated across all datasets, Hopx was consistently downregulated (Figure 1A and 1B), suggesting a role in transitions of astrocyte substates. We validated this downregulation by RNA sequencing of fluorescence-activated cell sorting (FACS)-isolated astrocytes from WT and 5×FAD mice (Figure 1C). Although recognized as a marker for quiescent neural stem cells in the dentate gyrus and reported in astrocytes¹², its function in astrocytes has not been documented.

**Figure 1.**
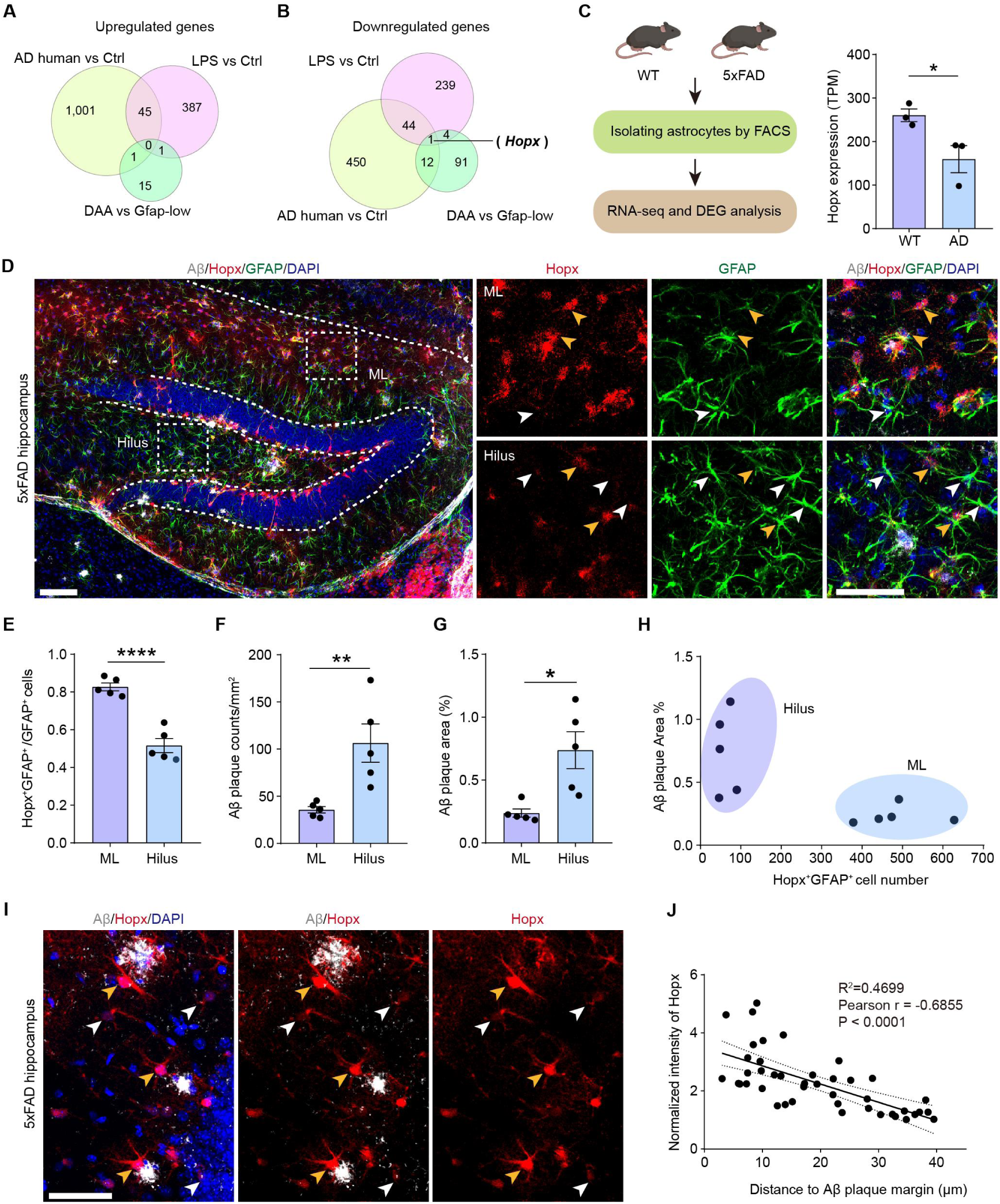
Identification of Hopx-expressing Astrocytes. (A-B) Overlapped astrocytic DEGs between human tissues (AD versus Ctrl), AD mice group (DAA versus Gfap-low) and inflammation mice (LPS treated versus Ctrl). (C) Astrocytes from WT and AD mice isolated by FACS and token RNA sequencing (left). Quantification of Hopx expression in astrocytes of WT and AD mice (right). Unpaired Student’s *t*-test. n=3 mice of each group. (D) Immunofluorescence staining of 5×FAD mouse hippocampus, Hopx (Red), Aβ (D54D2, gray), GFAP (green), DAPI (blue). Yellow arrows indicate astrocytes that expressed both Hopx and GFAP, white arrows indicate astrocytes that only expressed GFAP. ML: molecular layer. Scale bars, 100 μm (whole images) and 50 μm (boxed areas). (E-H) The proportion of Hopx-positive astrocytes in GFAP^+^ astrocytes (E), Aβ plaque counts (F) and Aβ plaque areas (G) in ML and Hilus. The correlation between Aβ plaque areas and the numbers of Hopx-positive astrocytes (H). Unpaired Student’s *t*-test. n=5 mice of each group. (I) Immunofluorescence staining of 5×FAD mouse hippocampus, Hopx (Red), Aβ (D54D2, gray), DAPI (blue) (I). White arrows indicate Hopx low expression cells, yellow arrows indicate Hopx high expression cells. Scale bars, 50 μm. (J) The correlation between Hopx expression in astrocytes and their distance to Aβ plaques. Pearson correlation analysis was performed to evaluate the relationship between the two variables, Pearson correlation coefficient (95% confidence interval: -0.8175 to -0.4850), n=43 cells from 3 mice. All data are presented as the mean ± SEM. *P < 0.05, **P < 0.01, ***P < 0.001. See also Figure S1

To characterize Hopx-expressing astrocytes, we employed immunofluorescence staining on brain sections from adult mice. We found that Hopx is expressed in a subset of astrocytes (Figure S1A). Upon quantification in hippocampus, we observed a higher density and proportion of Hopx-positive astrocytes in the hippocampal molecular layer (ML) compared to the hilus region (Figure S1B). This spatial distribution pattern was similarly evident in AD mouse models (Figure 1D). Interestingly, Aβ deposition was lower in the ML than in the hilus (Figure 1F-1H). Furthermore, Hopx expression negatively correlated with the distance to Aβ plaques (Figure 1I and 1J); astrocytes near plaques exhibited higher Hopx levels and often clustered around them (Figure S1C and S1D), implicating that Hopx-positive astrocytes respond to Aβ deposition.

### Hopx-positive astrocytes phagocytose Aβ and restrict Aβ deposition

As the deposition of Aβ is the pathological hallmark of AD^28, 29^, we studied the functional characteristics of Hopx-positive astrocytes in the context of Aβ phagocytosis. After tamoxifen injection in Hopx^CreER^;Ai9;5×FAD mice, Hopx-positive astrocytes were labeled with the fluorescent protein tdTomato. Flow cytometry analysis revealed a significantly higher proportion of Hopx-positive astrocytes engaged in Aβ phagocytosis compared to Hopx-negative astrocytes (Figure 2A and 2B), an effect more pronounced in 8-month-old 5×FAD mice (Figure S2A and S2B). Furthermore, we used serotype 5 adeno-associated virus (AAV5) combined with *GfaABC1D* promoter to label astrocytes in 5×FAD mice, and we found a greater percentage of Hopx-high astrocytes involved in phagocytosis compared to Hopx-low astrocytes (Figure 2C-2F). Thus, Hopx expression marks the astrocyte subpopulation with enhanced Aβ phagocytosis ability.

**Figure 2.**
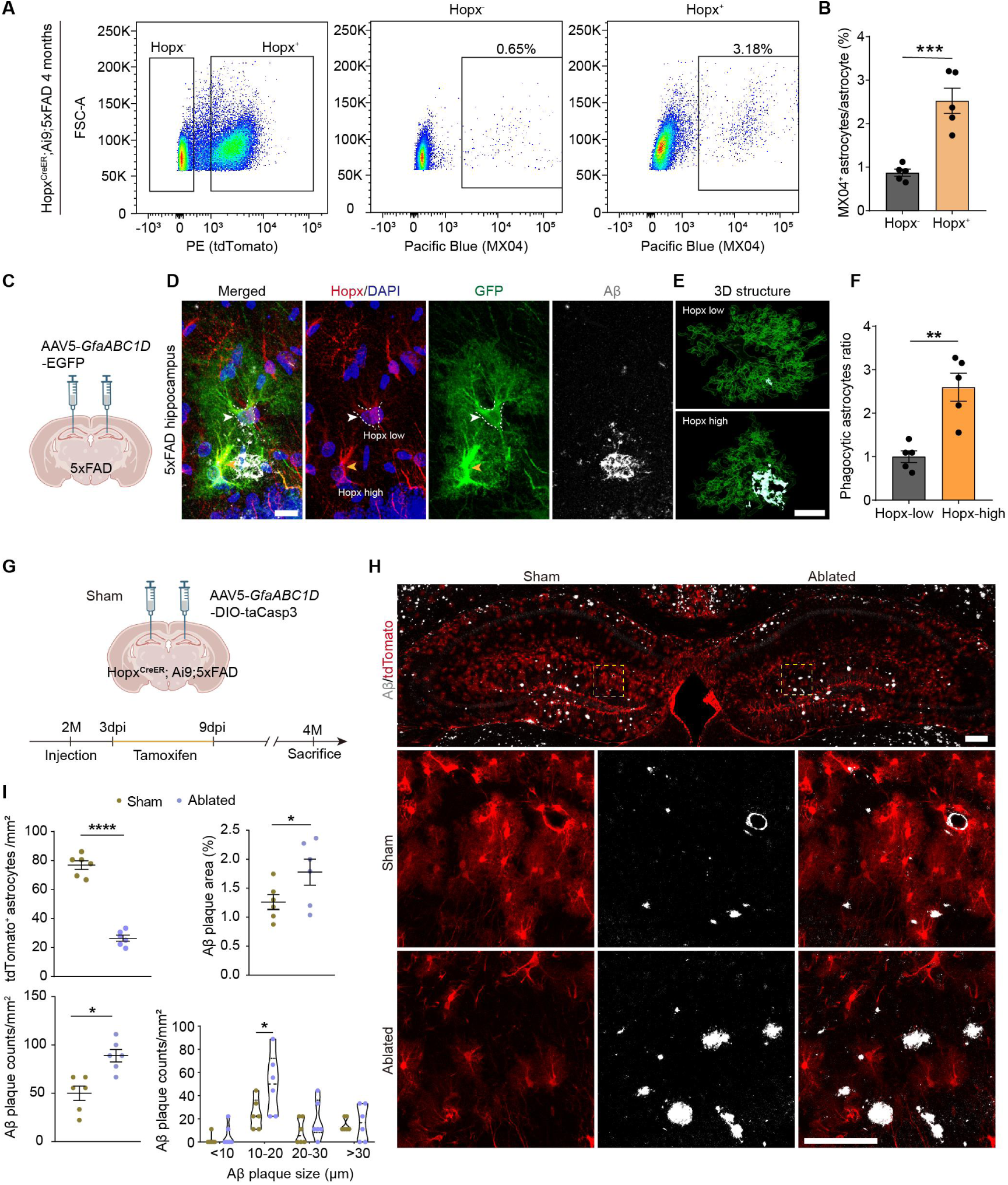
Hopx-positive astrocytes are essential in engulfment of Aβ plaque. (A) FACS analysis of astrocytes involved in engulfing Aβ plaque. PE-A channel was used to measure tdTomato signal, which marks Hopx-positive (Hopx^+^) astrocytes. Pacific Blue channel was used to measure MX04, a specific Aβ dye. Total astrocytes were marked by ACSA-2-APC. (B) Quantification for the percentage of cells engulfing Aβ inHopx^+^ and Hopx negative (Hopx^-^) astrocytes (B). Unpaired Student’s *t*-test, n= 5 mice of each group. (C) Schematic diagram showing the strategy of labelling astrocytes specifically with AAV5-*GfaABC1D*-GFP. (D) Immunofluorescence staining of AD mouse hippocampus. Hopx (Red), Aβ (D54D2, gray), DAPI (blue). White arrows show Hopx low expression (Hopx-low) cells, yellow arrows show Hopx high expression (Hopx-high) cells. Scale bar, 20 μm. (E) 3D structure of astrocytes and Aβ plaque. Scale bar, 20 μm. (F) Quantification for normalized ratio of phagocytic astrocytes (Aβ^+^EGFP^+^/GFP^+^). Unpaired Student’s *t*-test, n= 5 mice of each group. (G) Schematic procedure for AAV5-*GfaABC1D*-DIO-taCasp3 injection to ablate Hopx expression astrocytes. (H) Immunofluorescence staining of tdTomato (Red), Aβ (D54D2, gray). Scale bars, 200 μm (whole images) and 100 μm (boxed areas). (I) Quantitative analysis of tdTomato^+^ astrocytes, Aβ plaque area, Aβ plaque counts (unpaired Student’s *t*-test) and Aβ plaque counts of different size (two-way ANOVA). n= 6 mice of each group. All data are presented as the mean ± SEM. *P < 0.05, **P < 0.01, ***P < 0.001. See also Figure S2

To verify the role of Hopx-expressing astrocytes on Aβ deposition in AD mice, we injected the AAV5-*GfaABC1D*-DIO-taCasp3 into one side of the hippocampus of Hopx^CreER^;Ai9;5×FAD mice at 2 months of age and a control virus into the contralateral side (Figure 2G). Tamoxifen was administered to induce taCasp3 expression, thereby specifically ablate Hopx-positive astrocytes (Figure 2H). Seven weeks after tamoxifen injection, we observed a significant reduction of Hopx-positive astrocytes in the hippocampal DG region (Figure 2I). Analysis of Aβ plaque in areas where Hopx-positive astrocytes were ablated revealed a notable increase in Aβ deposition (Figure 2I). Furthermore, when examining plaques of varying sizes, we found an increase in the number of 10-20 µm Aβ plaques following the depletion of Hopx-positive astrocytes (Figure 2I). We also ablated Hopx-positive astrocytes at 2 months of age and examined Aβ deposition at 8 months of age (Figure S2C). We found that the number of Hopx-positive astrocytes remained significantly lower in the ablation group than in the control group (Figure S2D and S2E). Quantification of Aβ plaques revealed no significant difference in the number of plaques between the Hopx-positive astrocyte ablation group and the control group (Figure S2F and S2G). However, the area of the plaques was markedly increased, and the number of large plaques (> 500 μm²) was significantly elevated (Figure S2H and S2I). These results indicate that in AD mouse models, Hopx-positive astrocytes effectively reduce Aβ deposition through their enhanced ability to phagocytose Aβ.

### Astrocytic Hopx overexpression ameliorates Aβ pathology in 5×FAD Mice

Given the significant reduction of Hopx expression in astrocytes of 5×FAD mice (Figure 1A-C), we overexpressed Hopx in astrocytes and then assess the phagocytic capacity for Aβ. 5×FAD mice received AAV-mediated Hopx overexpression at 2 months of age and were analyzed 2 months later (Figure 3A). Flow cytometry showed an increased proportion of astrocytes engaged in Aβ engulfment upon Hopx overexpression (Figure 3B and 3C). Furthermore, immunofluorescence staining revealed a significant reduction in Aβ deposition in hippocampal (Figure 3D and 3E) and cortical (Figure S3A and S3B) regions overexpressing Hopx. These results indicate that Hopx plays a crucial role in Aβ phagocytosis of astrocytes.

**Figure 3.**
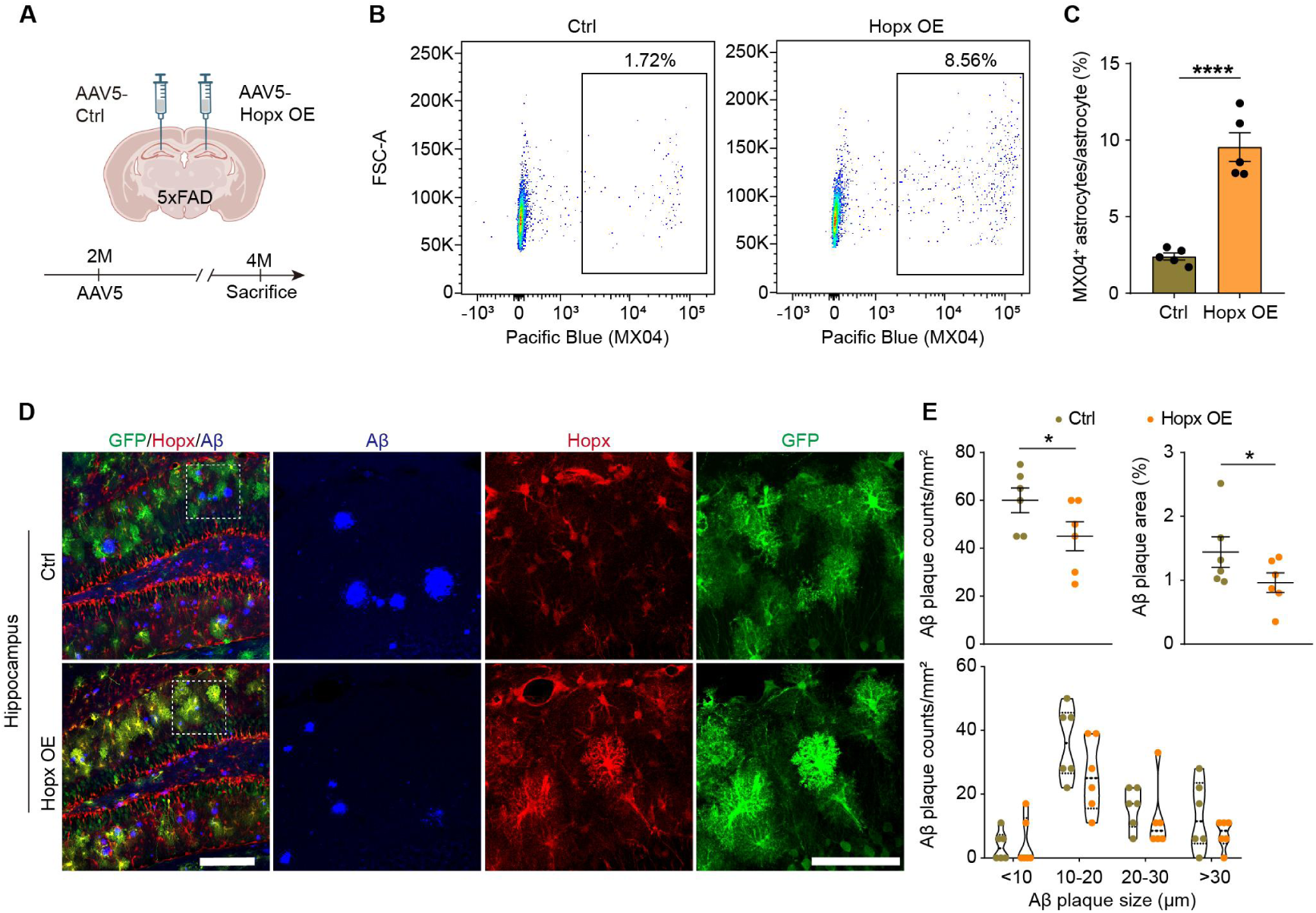
Astrocytic Hopx overexpression in 5×FAD mice reduces Aβ plaque deposit. (A) Schematic illustration of Hopx overexpression in 5×FAD mice. (B-C) Representative images of FACS analysis of astrocytes involved in engulfing Aβ plaque (B) and the quantification of the percentage of astrocytes engulfing Aβ in total astrocytes (C). Pacific Blue channel was used to measure the signal of MX04, a specific Aβ dye; astrocytes were marked by ACSA-2-APC. Unpaired Student’s *t*-test. n=5 mice of each group. (D-E) Immunofluorescence staining of 5×FAD mouse hippocampus, Hopx (red), GFP (green), D54D2 (gray), DAPI (blue). Scale bars, 200 μm (whole images) and 100 μm (boxed areas) (D) and the quantitative analysis of Aβ plaque area and Aβ plaque counts (unpaired Student’s *t*-test) and Aβ plaque counts of different size (two-way ANOVA) (E). n= 6 mice of each group. All data are presented as the mean ± SEM. *P < 0.05. See also Figure S3

### Loss of *Hopx* exacerbated Aβ plaque deposition

To further confirm the role of Hopx in astrocytes, we generated *Hopx*-knockout (KO) mice using CRISPR/Cas9 and crossed them with 5×FAD mice, and found that *Hopx* was efficiently knocked out (Figure 4A and S4A). Flow cytometry of astrocytes from the 5×FAD;KO mice showed a reduced proportion involved in Aβ engulfment (Figure 4B and 4C). Immunofluorescence analysis showed significantly increased Aβ plaque deposition in the hippocampus (Figure 4D and 4E) and cortex (Figure S4B and S4C) of these mice.

**Figure 4.**
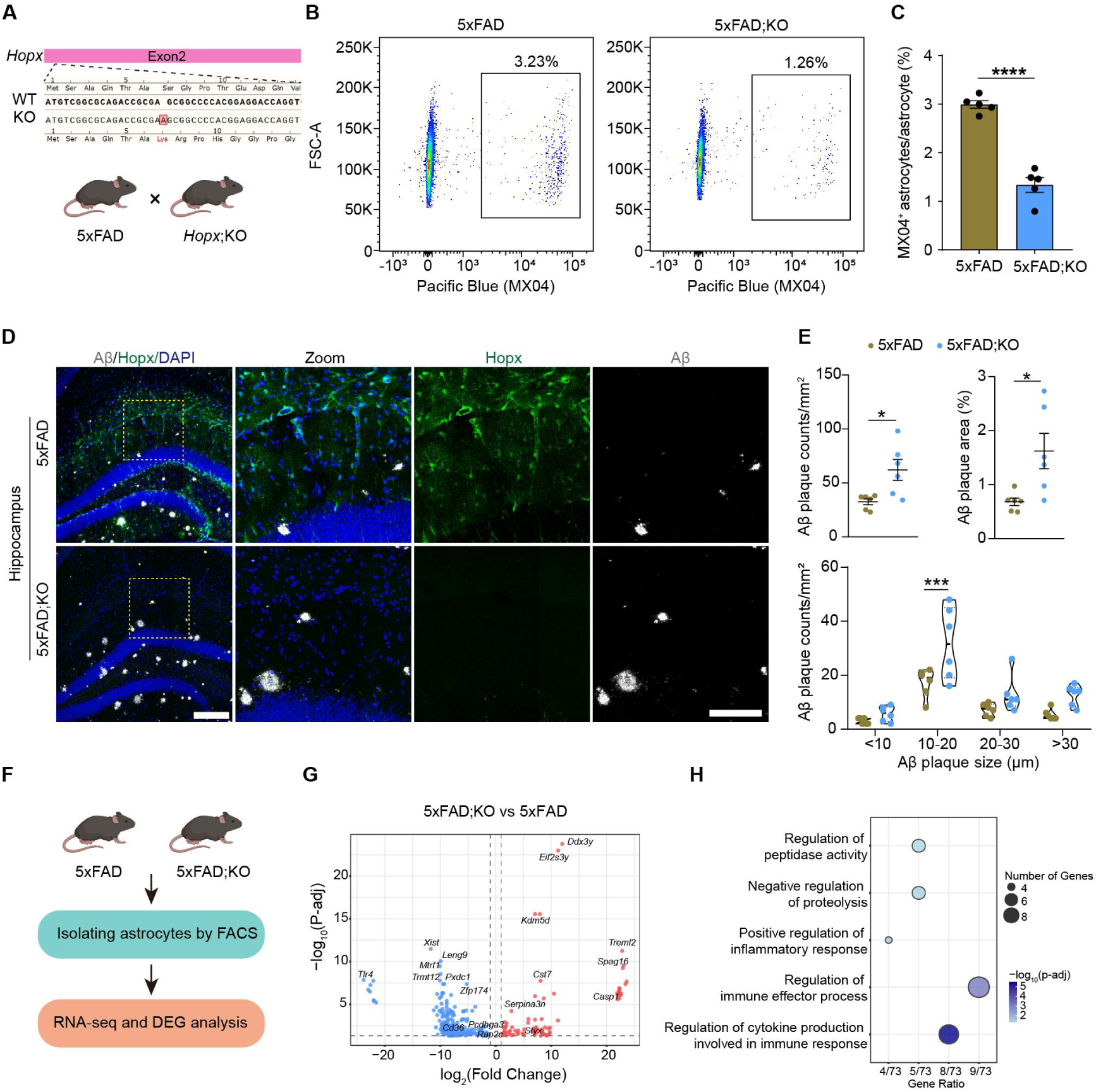
*Hopx* KO increases Aβ plaque deposition in 5×FAD mice. (A) The schematic diagram showing the construction of *Hopx* knockout mice, and mating strategy; (B) FACS analysis of astrocytes engulfing Aβ in 5×FAD and 5×FAD;KO mice. MX04 and ACSA-2-APC were used to mark Aβ and astrocytes, respectively. (C) The percentage of astrocytes engulfing Aβ in total astrocytes of 5×FAD and 5×FAD;KO mice. Unpaired Student’s *t*-test. n=5 mice of each group. (D) Immunofluorescence staining of 5×FAD and 5×FAD;KO mouse hippocampus, Hopx (green), D54D2 (gray), DAPI (blue). Scale bars, 200 μm (whole images) and 100 μm (boxed areas). (E) Quantitative analysis of Aβ plaque area and Aβ plaque counts (unpaired Student’s *t*-test) and Aβ plaque counts of different size (two-way ANOVA). n= 6 mice of each group. (F-H) The schematic diagram showing the strategy of astrocyte isolation and RNA sequencing (F), DEG volcano plots of astrocytes (G), and GO enrichment analysis (H). n= 3 mice of each group. All data are presented as the mean ± SEM. *P < 0.05, ***P < 0.001, ****P < 0.0001. See also Figure S4

To explore the underlying mechanisms by which Hopx regulates astrocytic phagocytosis of Aβ, we performed transcriptomic sequencing on FACS-isolated astrocytes from 5×FAD and 5×FAD;KO mice (Figure 4F). We found that a cohort of molecules related to phagocytosis (*Cd36, Elmo3* and *Mst1r*)^30^ and endocytic recycling pathway *(Egf, Pcdhga3, Rap2c, Pmel* and *Tlr4*)^31,32^ were downregulated in astrocytes of 5×FAD;KO mice (Figure 4G). Furthermore, in 5×FAD;KO mice, astrocytes highly expressed a series of proteolysis inhibitor molecules, such as *Styx*, *Cst7* and *Spint2*^33–35^ (Figure 4G). These findings suggest that Hopx, as a transcriptional regulatory molecule, modulates the capacity of astrocytes to phagocytose Aβ by regulating the expression of molecules associated with pathways including phagocytosis, endosomal recycling and proteolysis.

Interestingly, in 5×FAD;KO mice, astrocytes highly expressed *Serpina3n*, a molecule commonly used as a marker for reactive or toxic astrocytes in mouse models^9,15^ (Figure 4G). Furthermore, gene ontology (GO) enrichment analysis indicated that DEGs were associated with inflammatory responses, immune activation, and cytokine signaling pathways (Figure 4H). Collectively, these results suggest that Hopx deletion promotes astrocytic inflammatory responses and facilitates the transition toward a neurotoxic phenotype in AD mice.

### Hopx modulates astrocytic subpopulation transition in AD mice

Given that *Hopx* deletion promotes astrocytic inflammation and toxic property, we next investigated whether astrocyte-specific overexpression of Hopx reduces toxic astrocytes and preserves astrocytic homeostasis in the AD state. To verify the regulatory role of Hopx overexpression in the transition of distinct astrocyte subpopulations, we performed scRNA-seq following astrocyte-specific Hopx overexpression. Hippocampal tissues from WT-Ctrl, AD-Ctrl, and AD-Hopx mice were subjected to scRNA-seq (Figure 5A and 5B).

**Figure 5.**
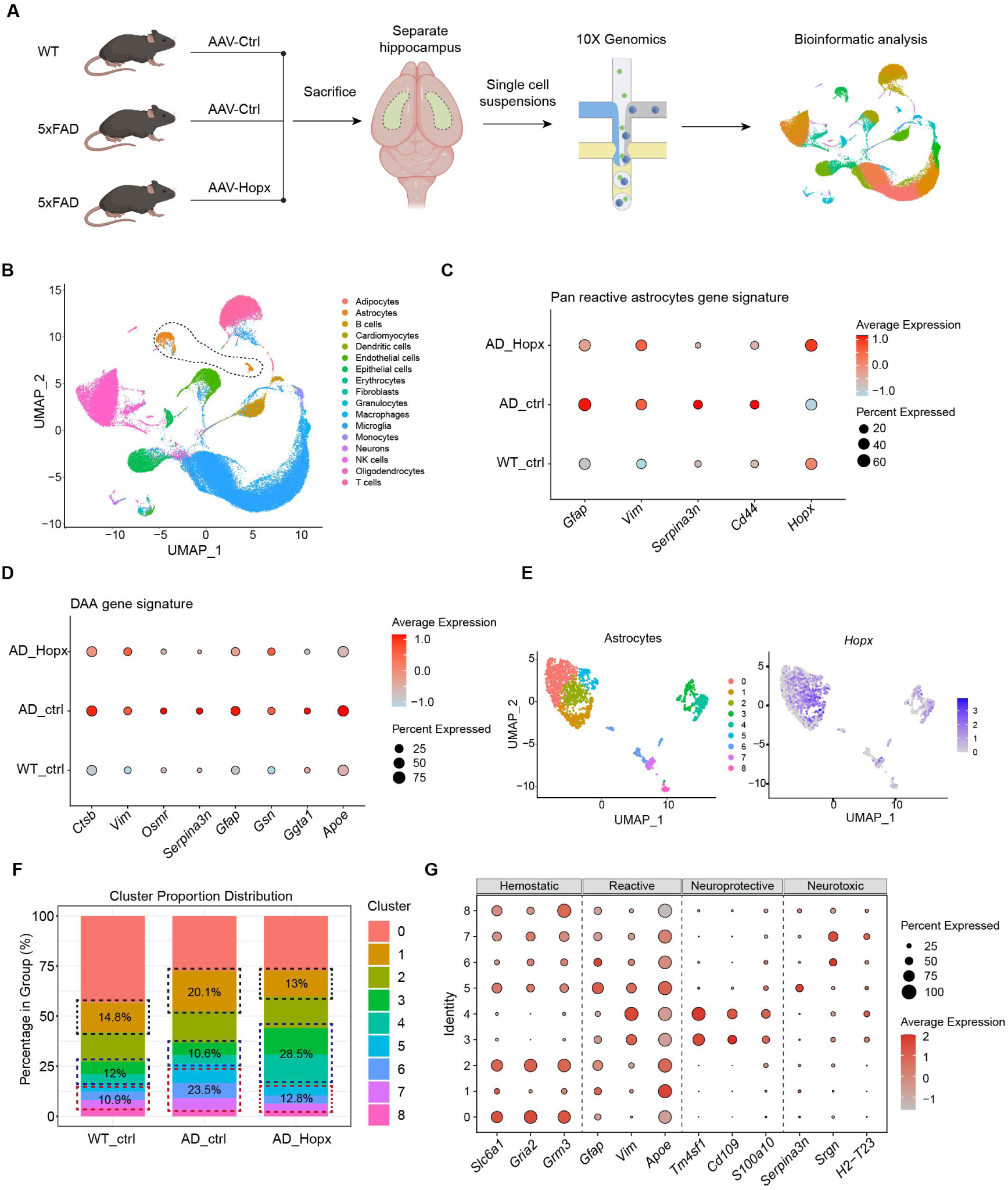
ScRNA-seq analysis of Hopx-overexpressing AD mice. (A) Schematic representation of the scRNA-seq experimental strategy employed in this study. (B) UMAP visualization of distinct cell types in adult hippocampal single-cell transcriptome data from the WT-Ctrl, AD-Ctrl, and AD-Hopx groups. (C) Expression of different genes corresponding to the pan-reactive astrocyte gene signature across the three groups. (D) Expression of different genes associated with the DAA gene signature across the three groups (E) UMAP visualization of astrocyte clusters (left) and Hopx expression across these distinct clusters (right). (F) The percentage of each astrocyte cluster (relative to total astrocytes) in each group. (G) Expression of different genes corresponding to hemostatic, neuroprotective, and neurotoxic gene signatures across distinct astrocyte clusters. See also Figure S5

Firstly, we compared the reactive state markers of astrocytes following Hopx overexpression. We observed that several pan-reactive astrocyte marker genes, such as *Gfap*, *Vim* and *Serpina3n* were upregulated in the AD-ctrl group but downregulated in the AD-Hopx group (Figure 5C). Meanwhile, the expression of certain disease-associated astrocytes (DAA) marker genes^18^ was also decreased after Hopx overexpression (Figure 5D). Previous studies have suggested that the transition of astrocytes into an activated state during inflammatory conditions and pathological states such as AD exhibits heterogeneity^8,14,36^. Specifically, some astrocytes adopted a neuroprotective phenotype, while others acquired neurotoxic characteristics. To investigate this heterogeneity, we compared the expression patterns of marker genes for both homeostatic and activated astrocytes across the three experimental groups. Our results revealed that Hopx overexpression led to downregulation of neurotoxic astrocyte marker genes and upregulation of neuroprotective astrocyte marker genes in the AD-Hopx group (Figure S5A). The homeostatic signature genes of astrocytes were significantly downregulated in AD-Ctrl group, while the expression levels of these homeostatic signature genes remained un-recovered even upon Hopx overexpression in astrocytes (Figure S5A). These findings indicate that, despite Hopx overexpression in astrocytes of AD mice, the cells still exhibited a prominent reactive state, yet this reactive state was characterized predominantly by neuroprotective astrocytic properties rather than neurotoxic phenotypes.

Furthermore, we performed a comprehensive analysis of astrocyte subsets. We found that nine distinct astrocyte subsets were present in all three experimental groups (Figure S5B). Moreover, *Hopx* was broadly expressed across all astrocyte subpopulations (Figure 5E and S5C), and Hopx overexpression altered astrocytic subset distribution (Figure 5F). The proportion of cluster 0 was decreased in AD-Ctrl and AD-Hopx mice. Cluster 1/5/6/7 astrocytes were increased in AD-Ctrl mice compared with WT-Ctrl mice, whereas this aberrant elevation was effectively normalized in AD-Hopx mice. In contrast, clusters 3 and 4 were expanded specifically in the AD-Hopx group (Figure 5F). Marker analysis identified cluster 0 as homeostatic (high *Slc6a1, Gria2, Grm3*), clusters 3/4 as neuroprotective (high *Tm4sf1, Cd109, S100a10*), and cluster 1 as reactive (high *Apoe, Gfap,* low *Slc6a1, Gria2, Grm3*), cluster 5/6/7 as neurotoxic (high *Serpina3n, Srgn, H2-T23*)^9,19^ (Figure 5G). These results indicate that Hopx regulates the transition of astrocyte subpopulations, promotes protective astrocytes and reduces neurotoxic astrocytes.

Notably, we observed that Sox9, recently reported to enhance astrocytic Aβ phagocytosis in AD mice^37^, was significantly upregulated upon Hopx overexpression (Figure S5D). Analysis of DEGs in Hopx-positive astrocytes from AD-Hopx versus AD-Ctrl mice revealed enrichment in pathways related to Aβ clearance and microtubule organization (Figure S5E and S5F), supporting a role for Hopx in promoting phagocytic function.

## DISCUSSION

Astrocytes, as the key cell type responsible for the clearance of Aβ plaques in the brain, exert a potent mitigating effect on AD progression via their phagocytic and scavenging functions^38, 39^. However, during the pathogenesis of AD, sustained stimulation by pathological factors such as inflammatory responses drive astrocytes to undergo a phenotypic switch from a homeostatic state to a reactive and even neurotoxic phenotype, and this phenotypic transition in turn accelerates the progression of AD^9,19,36,40^. In light of this, the exploration of intrinsic regulatory mechanisms within the AD brain that can simultaneously enhance the Aβ phagocytic capacity of astrocytes and preserve their homeostatic state holds important research value and significance. We identified *Hopx* as a molecule that is downregulated in astrocytes across inflammatory models, AD mouse models and the brain tissues of AD patients. Hopx is a non-DNA-binding homeodomain protein that acts as a regulatory scaffold: it exerts its functions by modulating the activity of transcription factors and chromatin-associated factors, thereby participating in the regulation of cardiac development^22^, whereas its functions in the nervous system remain largely elusive to date. Our findings revealed that Hopx-positive astrocytes act as “cleaners” in the brain of 5×FAD mice. Furthermore, Hopx modulates the Aβ phagocytic capacity of astrocytes, preserves astrocytic homeostasis and abrogates the generation of neurotoxic astrocytes. These results demonstrate the pivotal role of Hopx in astrocytes, rather than merely serving as a marker gene.

*Hopx* is well established as a marker gene for human outer radial glia during cortical development⁵. In mice, Hopx has been reported to be expressed in neural progenitors of the subventricular zone (SVZ) and the subgranular zone (SGZ) of the dentate gyrus, as well as in hippocampal astrocytes¹; however, the functional mechanisms of Hopx in these cell populations remain largely unclear. This study reveals the regulatory role of Hopx in astrocytes under the pathological state of AD. Notably, Hopx is still highly expressed in astrocytes of the brain under normal physiological conditions; therefore, its roles in regulating the normal physiological functions of astrocytes—such as the formation of the blood-brain barrier (BBB) and the regulation of signal communication between astrocytes and neurons — are all worthy of in-depth investigation. In addition, as a marker gene for neural stem cells in both the embryonic and adult stages, the regulatory function of Hopx in neurogenesis during the developmental stage and adult neurogenesis also holds significant value for further exploration.

Previous studies have shown that astrocyte-specific knockout of *Bace1* enhances Aβ plaque clearance in 5×FAD mice^41^, and astrocytic Sox9 overexpression promotes astrocytic phagocytosis of Aβ and preserves cognitive function in APP−NLGF mice^37^. Additionally, the transition of astrocytes from a homeostatic to a reactive phenotype is mediated by a class of specific molecular regulators, which sense pathological signals and drive transcriptional reprogramming, involving NF-κB, MAPK and mTOR signaling^40, 42–45^. However, few studies have identified molecular pathways that simultaneously modulate the Aβ phagocytic capacity and phenotypic states of astrocytes. In the present study, we found that knockout of *Hopx* led to a significant impairment in their ability to phagocytose Aβ plaques, consequently causing a marked accumulation of Aβ plaques in the mouse brain; meanwhile, the inflammatory signaling pathways in astrocytes were activated. In contrast, astrocyte-specific overexpression of Hopx potently enhanced their Aβ phagocytic capacity. ScRNA-seq analyses further confirmed that Hopx overexpression preserved the homeostatic phenotype of astrocytes and reduced the generation of neurotoxic astrocytes.

Mechanistically, Hopx acts as a regulatory scaffold to exert its biological functions by modulating the activity of transcription factors and chromatin-associated factors^46^. This characteristic suggests that Hopx is an upstream core regulator that exerts extensive control over the expression of downstream molecules, thereby achieving the dual regulation of astrocytic Aβ phagocytic capacity and transition of astrocytic subpopulation. Future studies are required to further identify the downstream transcription factors or chromatin-associated factors regulated by Hopx in astrocytes. Furthermore, our immunofluorescence staining results demonstrated that Hopx is not only localized in the nuclei of astrocytes but also widely expressed in the cytoplasm. This finding suggests that Hopx may possess non-nuclear functions independent of transcription factor regulation, and this potential mechanism of action is worthy of in-depth exploration.

In summary, our work delineates a potential therapeutic approach centered on Hopx-driven facilitation of astrocyte-mediated Aβ plaque phagocytosis and astrocyte substate transition in AD models. To further optimize these strategies for potential therapeutic translation, it will be important to extend these studies to a broader range of AD and neurodegenerative disease models and leverage additional molecular components of the astrocytic phagocytic machinery.

## Limitations of the study

Given the absence of aging as a crucial factor, the effects of astrocytic overexpression of Hopx on Aβ clearance and the amelioration of AD pathological phenotypes need to be further validated in additional AD mouse models and human-derived models, such as induced pluripotent stem cell (iPSC) systems or human brain organoid systems.

## ACKNOWLEDGMENTS

This work was partially supported by supported by National Key Research and Development Program of China (2024YFA1108000, 2021ZD0202500), National Natural Science Foundation of China (32130035), The Joint Project of the Yangtze River Delta Science and Technology Innovation Community (2024CSJZN0600), Changping Laboratory (2025B-07-15), and Lingang Labarotary (LGL-8998-11) and Shanghai Frontiers Science Center for Biomacromolecules and Precision Medicine at ShanghaiTech University. We thank the MultiOmics Core Facility, Molecular Imaging Core Facility, and Molecular and Cell Biology Core Facility at the School of Life Science and Technology, ShanghaiTech University, for providing technical support.

## AUTHOR CONTRIBUTIONS

D.D.C. and J.D. performed biochemical, morphology, histology, animal experiments and data analysis. X.Y.L., Y.Y.T. and Y.G. provided technical and data analysis assistance. J.D. and Z.G.L. designed the experiments and wrote the manuscript. Z.G.L. conceived the whole project.

## STAR★Methods

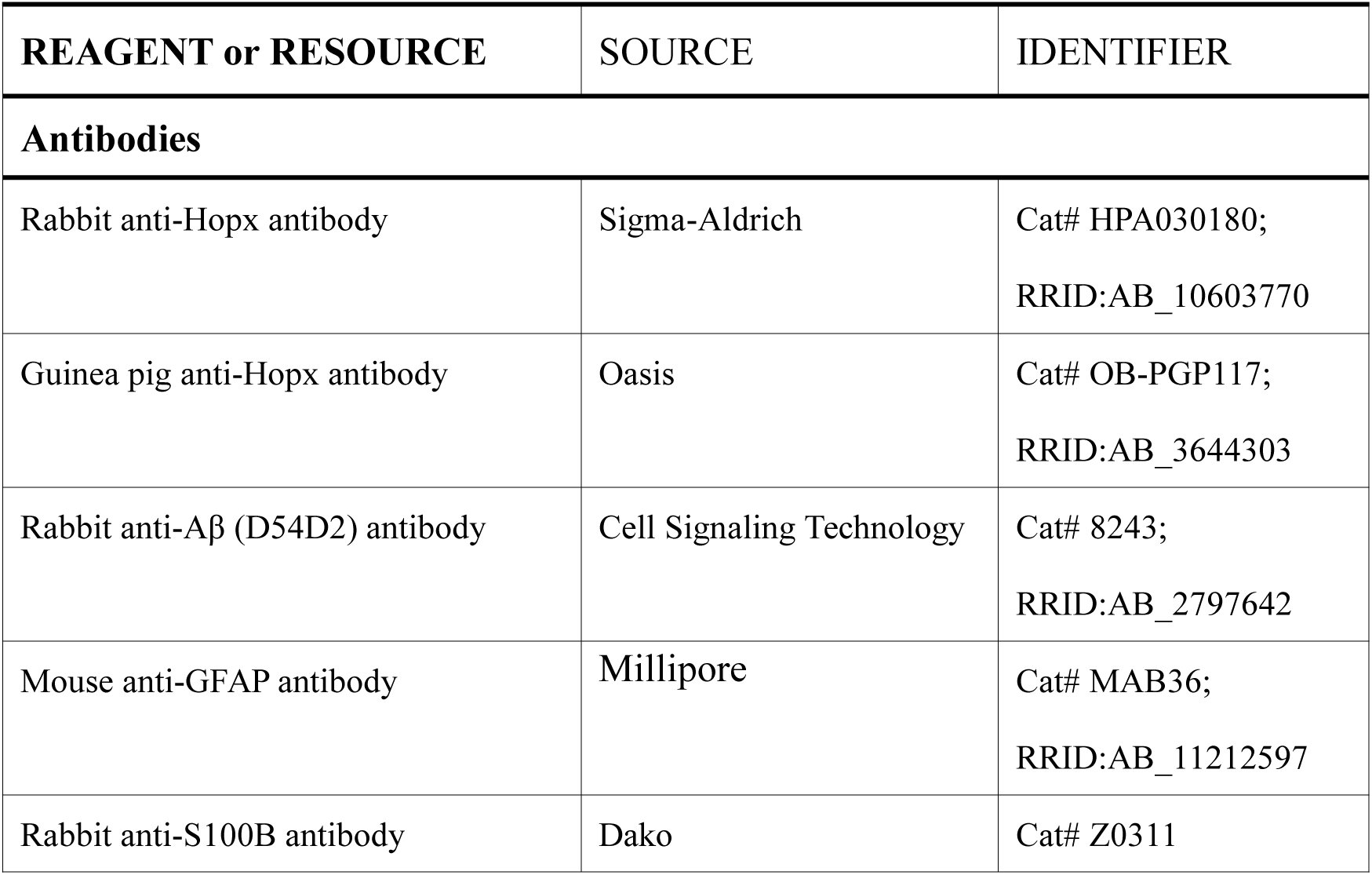
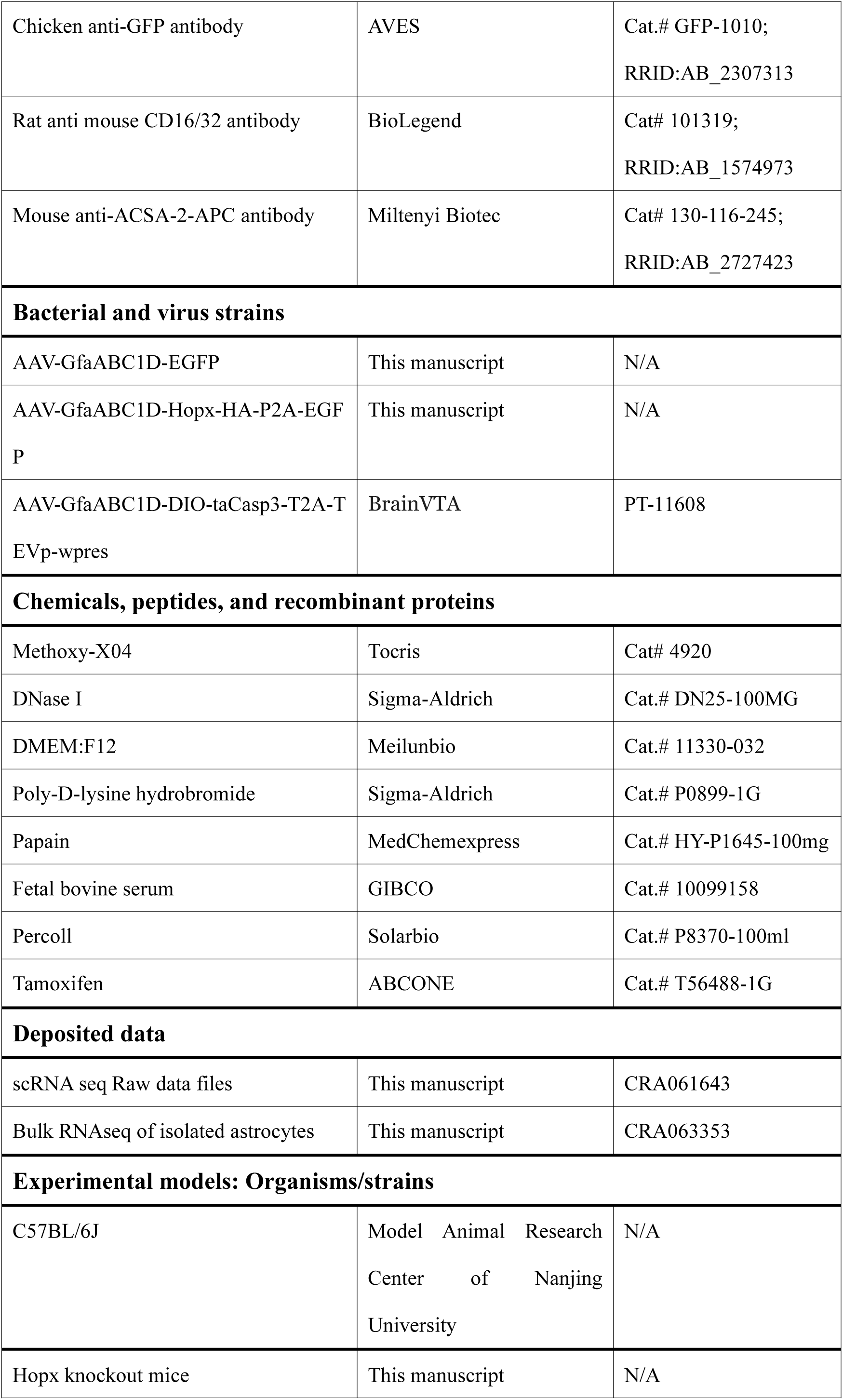
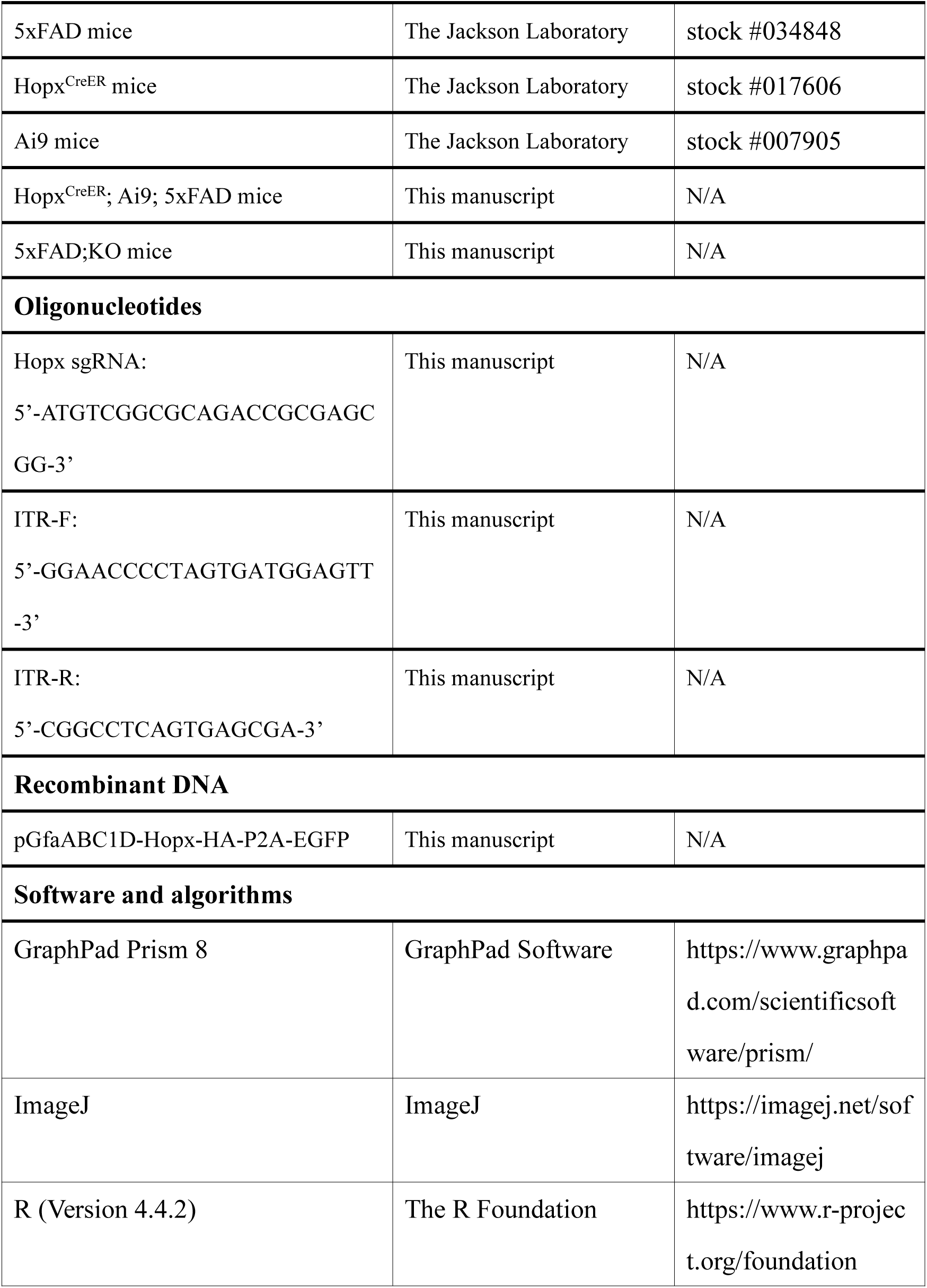
KEY RESOURCES TABLE.

## RESOURCE AVAILABILITY

### Lead Contact

Further information and requests for resources and reagents should be directed to and will be fulfilled by the Lead Contact, (Z.G.L.)

### Materials Availability

Requests for unique resources should be directed to the lead contact. A completed Materials Transfer Agreement may be required.

### Data and Code Availability

- The bulk RNA-seq and scRNA-seq raw sequencing data generated in this study, have been deposited and are publicly available at the Genome Sequence Archive under accession numbers CRA063353 and CRA061643, in the National Genomics Data Center database under project ID PRJCA057635.
- This paper does not report original code.
- The published article includes main datasets generated during this study. Any additional information is available from the Lead Contact upon request

## EXPERIMENTAL MODEL AND SUBJECT DETAILS

### Animals

The *5×FAD* (stock #034848), *Hopx^CreER^*(stock #017606), *Ai9*(stock #007905) and C57BL/6J(stock #000664) mice were purchased from the Jackson Laboratory. *Hopx* knock-out mice were generated by CRISPR-Cas9 technology with sgRNA sequence 5’-ATGTCGGCGCAGACCGCGAGCGG-3’. The *Hopx^CreER^*, *Ai9* and *5×FAD* mice were corssed to generate *Hopx^CreER^*;*Ai9*;*5×FAD* mice. The *Hopx*^-/-^ mice were mated with *5×FAD* mice to generate *5×FAD*;*Hopx^-/-^* (*5×FAD*;*KO*). Throughout the study, mice were housed under controlled temperature of 22°C and subjected to a 12-hour light/12-hour dark cycle with free access to water and food in animal facilities. All animal were performed with procedures strictly adhered to the ethical guidelines set forth by the Institutional Animal Care and Use Committee of ShanghaiTech University.

## METHOD DETAILS

### Plasmid construction

For Hopx overexpression, *Hopx-HA* fragments was synthesized and cloned into an AAV vector with the *GfaABC1D* promoter.

### Flow cytometry analysis of Aβ phagocytosis

The in vivo phagocytosis assay was modificated based on previously reported methods^1^. Hopx^CreER^;Ai9;5×FAD mice, 5×FAD;KO mice and 5×FAD mice at 4 months of age were intraperitoneally injected with 10 mg/kg methoxy-X04. After anesthesia, mice were perfused with ice-cold PBS and isolated cortex and hippocampus quickly. The isolated tissue were cut into pieces, dissociated enzymatically in Papain (1.5 mg/mL, MCE), DNase I (50 U/mL, Sigma) in DMEM F/12 (Meilunbio) for 30 min at 37℃. Homogenize the dissociated hippocampi by pipetting gently up and down. The homogenates were filtered through a 70 μm cell strainer and centrifuged at 300g for 5 min. Then, cells were resuspended in 30% Percoll (Solarbio) in ice-cold HBSS and centrifuged at 700 g at 4℃ for 10 min. The cell pellet was washed two times by ice-cold DPBS and after centrifugation and rewash, then resuspended in FACS buffer (0.5% BSA in DPBS) with CD16/CD32 (BioLegend) at 4℃ for 10 min. Then cells were incubated with ACSA2-APC (Miltenyi) in FACS buffer for 45 min at 4℃. Then cells were token for flow cytometry analysis (BD Fortessa).

### Tamoxifen administration

Stock solutions of tamoxifen (ABCONE) were prepared at concentration of 20 mg/ml in corn oil (ABCONE). 3 days after virus injection, *Hopx^CreER^;Ai9;5×FAD* mice were injected with tamoxifen intraperitoneally at dosage of 100 mg/kg body weight for 1week.

### Immunofluorescence

After anesthesia, mice were perfused with ice-cold PBS followed by 4% paraformaldehyde (PFA), and brain tissues were fixed overnight in 4% PFA, Brain sections were cut at 50 μm thickness coronally by using vibratome. For Immunofluorescence, brain sections were incubated with solution containing 0.3% Triton X100 and 10% BSA in PBS for 2 hours to block nonspecific binding. The primary antibodies were diluted in a solution of 0.3% Triton X100 and 2%BSA in PBS, for overnight at 4℃. Then, followed by washes, the samples were incubated with appropriate secondary antibodies labeled by Alexa Fluor^TM^ 488, 555, or 647 (diluted at 1:1000) for 2 hours at room temperature. Cell nuclei were counter-stained using Diamidino-2-Phenylindole, Dihydrochloride (DAPI, Beyotime Biotechnology). Images were collected using an Olympus FV3000 confocal microscope.

### Preparation of AAV

AAV was generated in HEK293T cells via triple transfection with the AAV helper plasmid (phelper), pAA2/5 (Addgene #104964), and the target plasmid carrying the coding sequence of EGFP or Hopx. Polyethyleneimine (PEI 40000, Yeasen, #40816ES02) served as the transfection reagent for delivering plasmids into HEK293 cells, with the mass ratio of PEI to total plasmid DNA set at 3:1. Viral particles were purified by iodixanol gradient ultracentrifugation (Sigma-Aldrich, #D1556-250ML) conducted at 48,000×g and 4℃ for 2.5 hours, then concentrated in phosphate-buffered saline (PBS) using Amicon Ultra-100 centrifugal filter units (Millipore, #UFC910096). The purified virus was aliquoted into single-use fractions and stored at -80℃ until subsequent in vivo injection. Viral titer was quantified as virus genomes per milliliter by detecting the levels of inverted terminal repeat (ITR) through quantitative real-time polymerase chain reaction (qPCR). Prior to the release of viral DNA from viral particles, all exogenous free DNA was eliminated by digestion with DNase I (Thermo Fisher Scientific, #EN0525). The qPCR assay was performed with SYBR Green dye (Bimake, #B21702), utilizing the following ITR-specific primers: 5’-GGAACCCCTAGTGATGGAGTT-3’ (forward) and 5’-CGGCCTCAGTGAGCGA-3’ (reverse).

### Stereotaxic injection

In brief, mice were anesthetized with intraperitoneal injection of 2,2,2-tribromoethanol (Sigma-Aldrich). The head was disinfected with 70% ethanol, followed by a skin incision. All AAVs were injected at 1E+10 vg in 0.5 μL, with three sets of coordinates applied: anterior-posterior (AP) -2.0 mm relative to the bregma, medial-lateral (ML) ±1.5 to 1.7mm, and dorsal-ventral (DV) -2.0 mm (for the hippocampus) or -1.0 mm (for the cortex) relative to the brain surface. After injection, the incision was closed with a silk suture. Injected mice were allowed to recover in a heated cage for 1 hour before being returned to their home cages.

### Single-cell RNA sequencing and data analysis

After anesthesia, blood was collected from the mouse heart using a 1ml syringe, and hippocampal tissues were harvested from at least two mice per group (WT-Ctrl, AD-Ctrl, and AD-Hopx). All procedures ranging from tissue digestion to scRNA-seq library preparation and sequencing were performed by Novogene Bioinformatics Technology Co., Ltd. (Tianjin, China).

Single-cell suspensions were sequentially filtered through 70-μm and 30-μm cell strainers (Miltenyi Biotech, Part Numbers: 130-098-458 and 130-098-462) prior to treatment with Red Blood Cell Lysis Solution (Miltenyi Biotech, Part Number: 130-094-183). Cell viability was assessed using the Countstar Rigel instrument (Alit Biotech), and dead cell depletion was performed based on the viability results using the Dead Cell Removal Kit (Miltenyi Biotech, Part Number: 130-090-101).Ultimately, the cells were resuspended in 1× PBS (Invitrogen) containing 0.04% BSA to a final concentration of 700–1200 cells/μL, followed by processing with the 10x Chromium Single Cell 3’ Kit (v4, Part Number: 1000691). The cell suspension was loaded onto a 10x Chromium Chip (v4, Part Number: 1000690) and barcoded using a 10x Chromium Controller. RNA from the barcoded cells was then reverse-transcribed, amplified, and converted into sequencing libraries using the 10x Library Construction Kit (v4, Part Number: 1000694) in accordance with the manufacturer’s protocols. Sequencing was carried out on an Illumina NovaSeq platform using 150-bp paired-end reads.

Fastp software (v0.20.0) was utilized to conduct basic quality control statistics on the raw sequencing reads. Typically, 10x Genomics® Cell Ranger software accepts FASTQ files directly generated from raw base call (BCL) files by Illumina sequencers as input. Raw reads (or cleaned reads if preprocessed) were demultiplexed and aligned to the reference genome via the 10X Genomics Cell Ranger pipeline (https://support.10xgenomics.com/single-cell-gene-expression/software/pipelines/late st/what-is-cell-ranger) using the ‘cellranger count’ function with default parameters. Cells with fewer than 2,000 unique molecular identifiers (UMIs) were excluded from further analysis.

In R software (version 4.4.2), datasets from the WT-Ctrl, AD-Ctrl, and AD-Hopx groups were integrated into a single Seurat object (using Seurat v5)^2^ with filtering thresholds set as 200–6,000 nFeature_RNA and <25% mitochondrial gene percentage (percent.mt). Subsequent analyses included data normalization, identification of variable features (FindVariableFeatures), data scaling (ScaleData), principal component analysis (PCA), uniform manifold approximation and projection (UMAP), clustering (FindClusters), and cell type annotation using the SingleR (v2.8.0). Astrocyte clusters were subset for subcluster analysis with a resolution of 0.5. Gene Ontology (GO) enrichment analysis of the identified marker gene sets was conducted using the clusterProfiler (v4.14.3 R package) and org.Mm.eg.db (v3.20.0 R package).

### Bulk RNA-seq and analysis

Astrocytes were harvested from 5×FAD and 5×FAD;Hopx knockout (KO) mice, and flash-frozen in liquid nitrogen immediately after the addition of Trizol reagent. RNA-seq library preparation and sequencing were conducted by Shanghai Majorbio Bio-pharm Technology Co., Ltd. For transcriptome library construction, the SMART-Seq_V4 Ultra Low Input RNA Kit for Sequencing (Clontech Laboratories, San Diego, CA, USA) was utilized, with 10 ng of total RNA as the input material. Following quantification using a Qubit 4.0 Fluorometer, the paired-end RNA-seq libraries were subjected to sequencing on a NovaSeq X Plus sequencer (Illumina), with a read length of 2 × 150 bp. Raw paired-end reads underwent trimming and quality control using fastp software with default parameters to obtain high-quality clean reads. Subsequently, the clean reads from each sample were individually aligned to the mouse reference genome (GRCm38/mm10) in stranded mode via the HISAT2 alignment tool. The mapped reads of each sample were then assembled using StringTie software. To quantify gene expression levels, the transcripts per million reads (TPM) method was employed to calculate the expression abundance of each transcript. Differential expression analysis between the two experimental groups was performed using the DESeq2 R package to identify DEGs.

## QUANTIFICATION AND STATISTICAL ANALYSIS

Statistical analyses were conducted using GraphPad Prism 8 software. All results are expressed as mean ±SEM. For statistical evaluation of differences: unpaired student’s *t*-test was applied for comparisons between two groups, while two-way ANOVA followed by Sidak’s multiple comparisons test was used to assess multiple comparisons among three or more groups.

