## Supplemental information for "Hopx expression marks Aβ clearance astrocytes in Alzheimer’s disease"

**SUMMERY**

Astrocytic dysfunction is closely associated with nearly all types of neurological diseases. Targeting astrocyte regulation has thus emerged as an important potential therapeutic strategy for neurological disorders. However, mechanisms that induce astrocyte dysfunction under pathological conditions have not been fully elucidated. In this study, we identified that Homeodomain-only protein X (*Hopx*) is downregulated in astrocytes across inflammatory models, Alzheimer's disease (AD) mouse models, and brain tissues from AD patients. In the brains of AD mice, Hopx-positive astrocytes exhibit efficient β-amyloid (Aβ) plaque clearance, while knockout of *Hopx* leads to astrocytic dysfunction. Conversely, astrocyte-specific overexpression of Hopx not only significantly enhances their Aβ phagocytic function but also effectively reduces the generation of neurotoxic astrocytes while increasing protective astrocytes. In summary, this study demonstrates the core regulatory mechanism underlying astrocytic dysfunction under AD pathological conditions and provides important potential targets for developing therapeutic strategies for AD by targeting astrocyte regulatory pathways.

### Supplemental information titles and legends

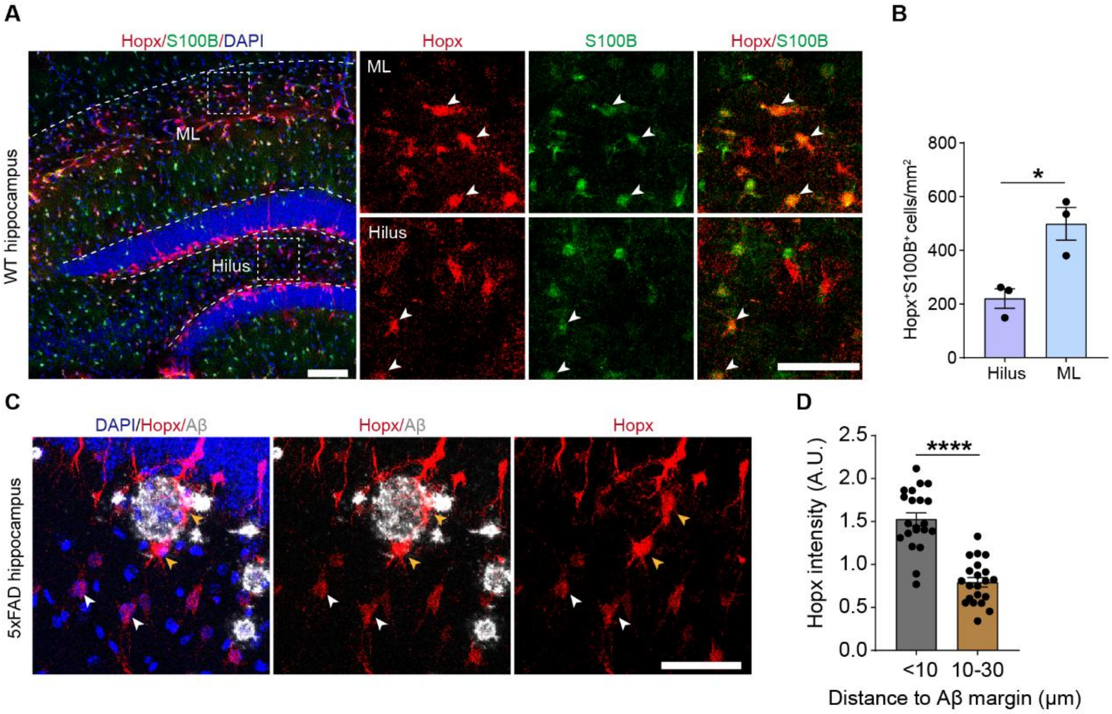


**Figure S1. Hopx expression pattern in WT and 5×FAD mice.**

(A) Representative images showing the expression of Hopx in hippocampus of WT mice, Hopx (red), S100B (green), and DAPI (blue). Scale bars, 100 μm (whole images) and 50 μm (boxed areas).

(B) Quantitative analysis of $Hopx^{+}S100B^{+}$ astrocytes in molecular layer (ML) and Hilus of DG. Unpaired Student's *t*-test. n=3 mice of each group.

(C) Representative images showing the expression of Hopx in hippocampus of 5×FAD mice, Hopx (red), Aβ (D54D2, gray), DAPI (blue). White arrows indicate Hopx-low cells, yellow arrows indicate Hopx-high cells. Scale bars, 50 μm.

(D) The intensity of Hopx signals in astrocytes at areas with indicated distance to Aβ plaque margins. Unpaired Student's *t*-test. n=3 mice.

Data were presented as mean ± SEM. * $P < 0.05$; ****$P < 0.0001$.

See also Figure 1.

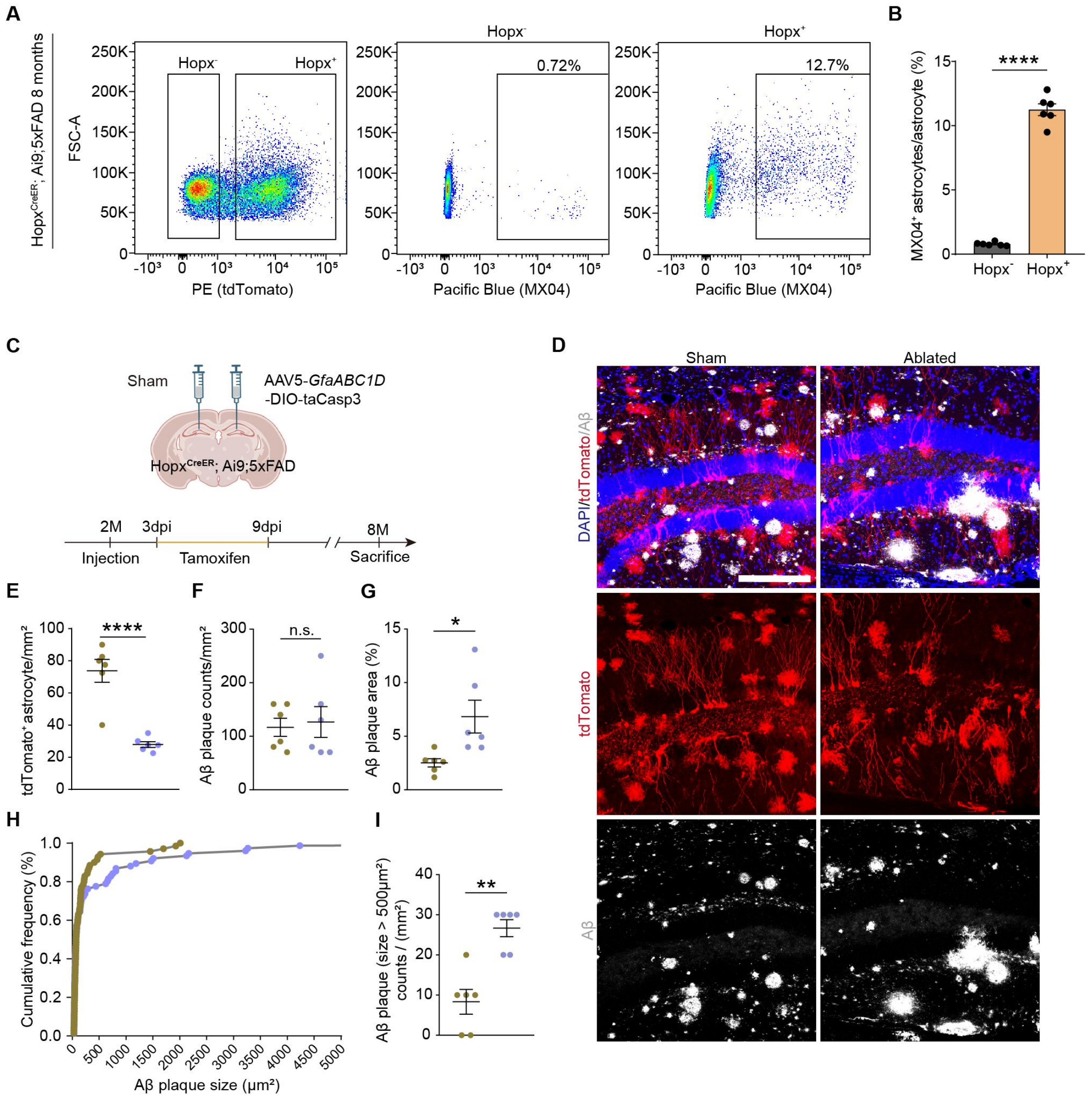


**Figure S2. Hopx-positive astrocytes engaging in Aβ phagocytosis in 8-month-old 5×FAD mice.**

(A) FACS analysis of astrocytes involved in engulfing Aβ. PE-A channel was used to measure tdTamoto signal, which marks Hopx$^{+}$ astrocytes, Pacific Blue channel was used to measure MX04 signal, which marks Aβ, and ACSA-2-APC was used to label total astrocytes.

(B) Quantification for the percentage of cells engulfing Aβ in Hopx$^{+}$ and Hopx$^{-}$ astrocytes. Unpaired Student's *t*-test. n= 6 mice of each group.

Data are presented as mean ± SEM. ****$P < 0.0001$.

(C) Schematic procedure for AAV5-GfaABC1D-DIO-taCasp3 injection to ablate Hopx expression astrocytes.

(D) Immunofluorescence staining of DAPI (blue), tdTomato (Red), Aβ (D54D2, gray).

Scale bars, 200 μm.

(E-G) Quantitative analysis of tdTomato+ astrocytes (E), Aβ plaque counts (F), Aβ plaque area (G), (unpaired Student's t-test). n= 6 mice of each group.

(H-I) Cumulative frequency of Aβ plaques (H), quantitative analysis of Aβ plaques > 500 $\mu m^2$ (student's *t*-test). n= 6 mice of each group.

All data are presented as the mean ± SEM. *P < 0.05, **P < 0.01, ***P < 0.001. None sense (n.s.)

See also Fig. 2.

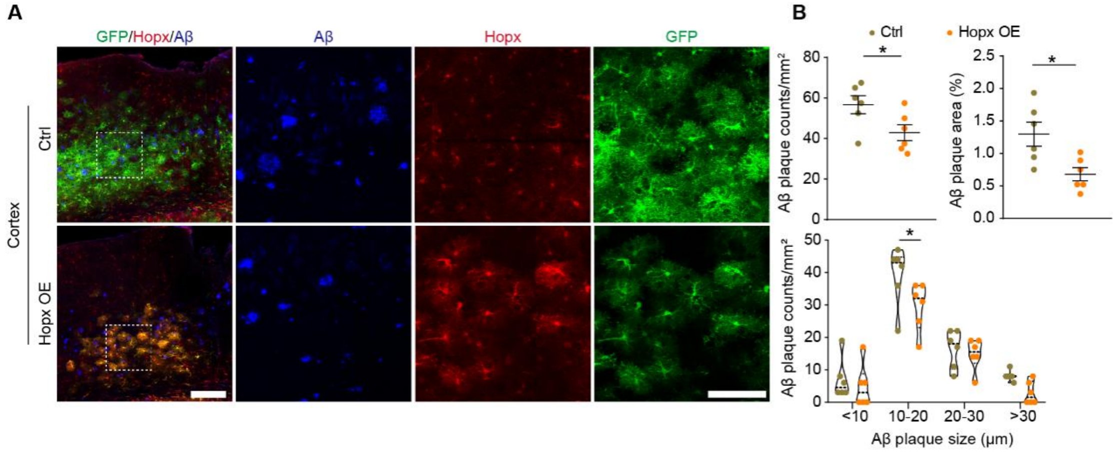


**Figure S3. Hopx overexpression reduces Aβ plaque deposit in cortex of 5×FAD mice.**

(A) Immunofluorescence staining of 5×FAD mouse cortex, Hopx (red), GFP (green), D54D2 (gray), DAPI (blue). Scale bars, 200 μm (whole images) and 100 μm (boxed areas).

(B) Quantitative analysis of Aβ plaque area and Aβ plaque counts (unpaired Student's *t*-test), and Aβ plaque counts of different size (two-way ANOVA). n=6 mice of each group.

Data are presented as mean ± SEM. *$P < 0.05$.

See also Fig. 3.

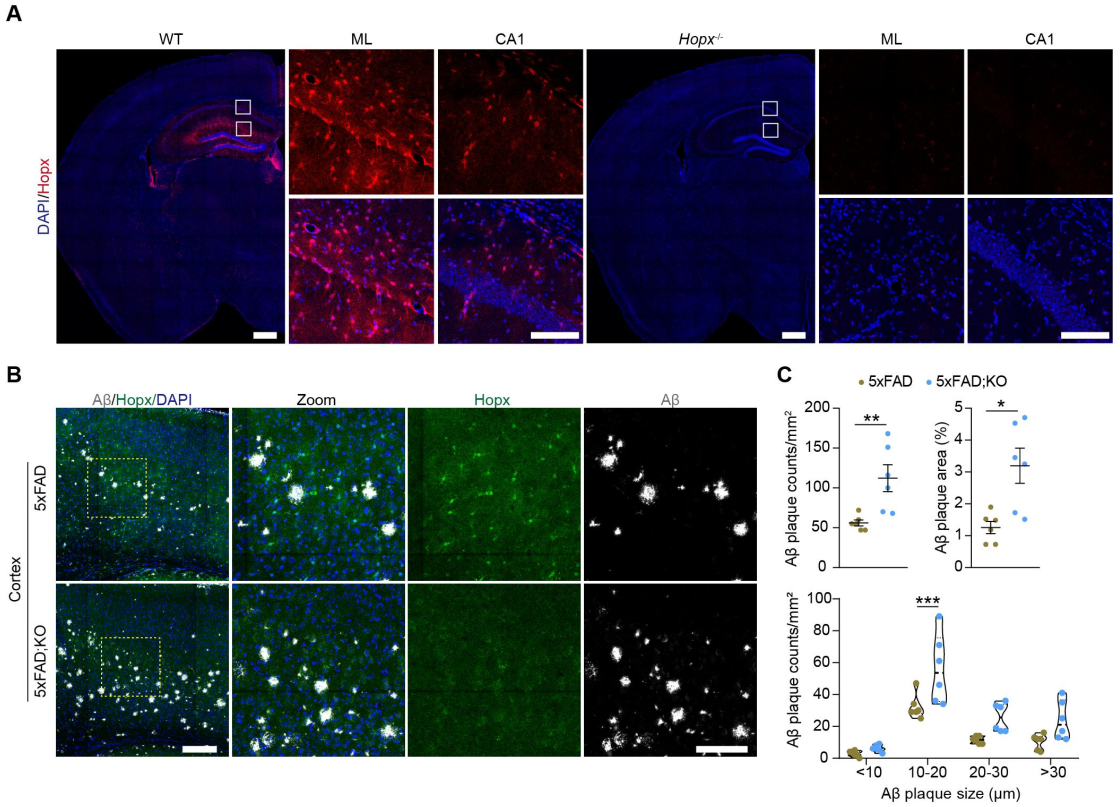


**Figure S4.** ***Hopx*** **KO increases cortical Aβ plaque deposition in 5×FAD mice.**

(A) Immunofluorescence staining of adult WT and Hopx$^{-/-}$ mice, Hopx (red), DAPI (blue), Scale bars, 500 μm (whole images) and 100 μm (boxed areas).

(B) Immunofluorescence staining of 5×FAD and 5×FAD;KO mouse cortex, Hopx (green), D54D2 (gray), DAPI (blue), Scale bars, 200 μm (whole images) and 100 μm (boxed areas) .

(C) Quantitative analysis of Aβ plaque aera and Aβ plaque counts, and Aβ plaque counts of different size. Unpaired Student's *t*-test and two-way ANOVA. n=6 mice of each group.

Data are presented as mean ± SEM. *$P$ < 0.05, ***$P$ < 0.001.

See also Fig. 4.

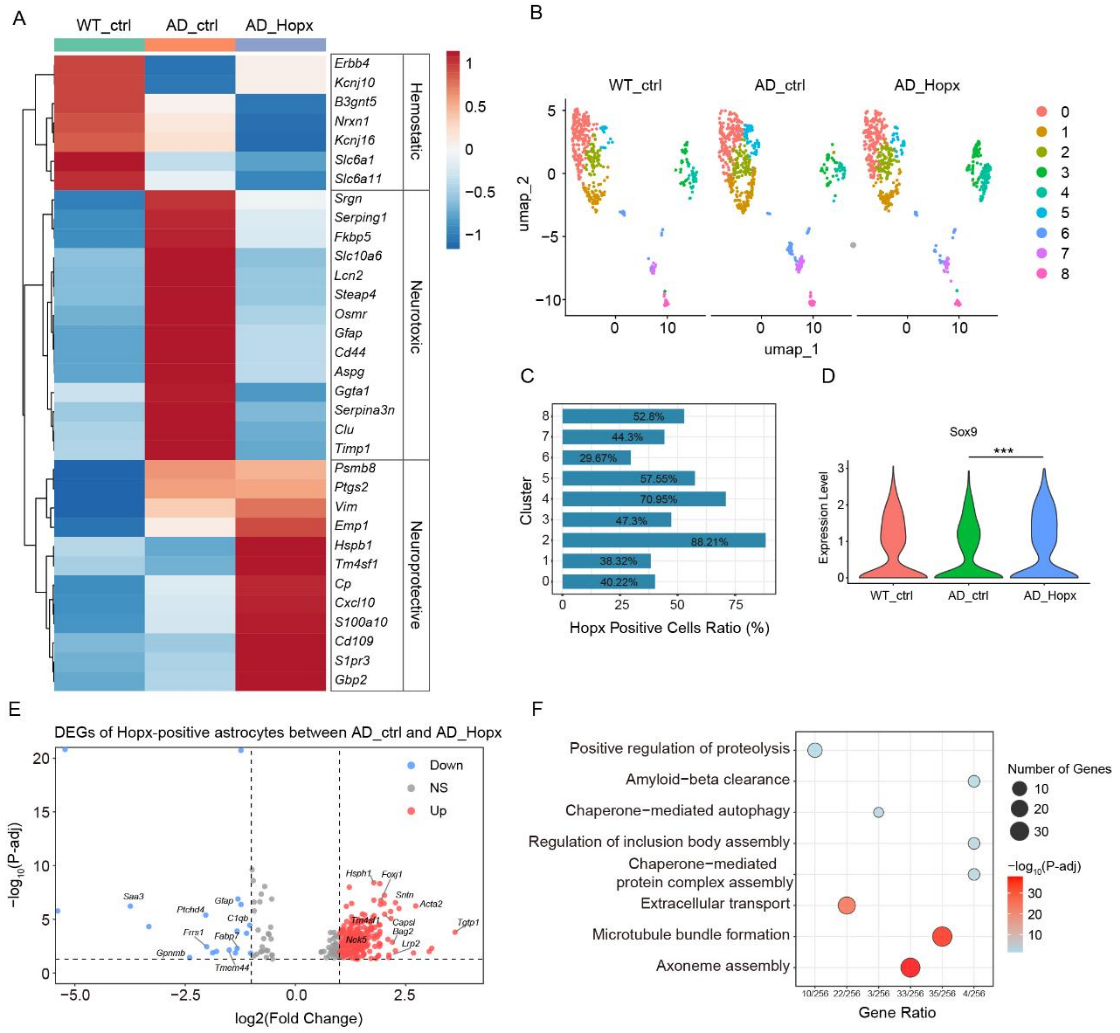


**Figure S5. Hopx overexpression modulates the phenotypic states of astrocytes.**

(A) Heatmap of hemostatic, neurotoxic and neuroprotective astrocytes marker genes in indicated groups.

(B) UMAP analysis of astrocytes subclusters in indicated groups.

(C) The ratio of $Hopx^+$ astrocytes in each subcluster.

(D) Violin plot of Sox9 expression in astrocytes of indicated groups.

(E) Volcano plot showing DEGs of $Hopx^+$ astrocytes between AD-Hopx versus AD-Ctrl group.

(F) GO analysis of up-regulated DEGs of Hopx positive astrocytes between AD-Hopx versus AD-Ctrl group.
